# Preclinical evaluation of tissue-selective gene therapies for congenital generalised lipodystrophy

**DOI:** 10.1101/2024.06.27.600757

**Authors:** Mansi Tiwari, Ahlima Roumane, Nadine Sommer, Weiping Han, Mirela Delibegović, Justin J. Rochford, George D. Mcilroy

## Abstract

Lipodystrophy is a rare disorder which can be life-threatening. Here individuals fail to develop or maintain appropriate adipose tissue stores. This typically causes severe metabolic complications, including hepatic steatosis and lipoatrophic diabetes. There is no cure for lipodystrophy, and treatment options remain very limited. Here we evaluate whether tissue-selective adeno-associated virus (AAV) vectors can provide a targeted form of gene therapy for lipodystrophy, using a preclinical lipodystrophic mouse model of *Bscl2* deficiency. We designed AAV vectors containing the mini/aP2 or thyroxine-binding globulin promoter to selectively target adipose or liver respectively. The AAV-aP2 vectors also contained the liver-specific microRNA-122 target sequence, restricting hepatic transgene expression. Systemic delivery of AAV-aP2 vectors overexpressing human *BSCL2* restored adipose tissue development and metabolic health in lipodystrophic mice without detectable expression in the liver. High doses (1×10^12^ GCs) of liver-selective vectors led to off target expression and adipose tissue development, whilst low doses (1×10^10^ GCs) expressed selectively and robustly in the liver but did not improve metabolic health. This reveals that adipose tissue-selective, but not liver directed, AAV-mediated gene therapy is sufficient to substantially recover metabolic health in generalised lipodystrophy. This provides an exciting potential new avenue for an effective, targeted, and thereby safer therapeutic intervention.

## INTRODUCTION

Gene therapy aims to provide therapeutic benefit to individuals suffering from diseases or disorders that currently have no cure. This can be achieved using various strategies, including gene replacement, gene silencing or gene editing technologies ^1^. Whilst numerous delivery platforms are available for gene therapy, adeno-associated virus (AAV) vectors appear to be the safest and most effective vehicle for therapeutic purposes ^2, 3^. A recent meta-analysis of two decades of AAV usage in clinical settings identified 136 unique clinical trials with AAV products designed to treat 55 disorders ^3^. These intensive research efforts have led to multiple AAV based gene therapies (Glybera (discontinued 2017), Beqvez, Elevidys, Hemgenix, Luxturna, Roctavian, Upstaza and Zolgemsa) successfully gaining regulatory approval for the treatment of a variety of disorders.

AAV-mediated gene therapies are particularly effective to treat rare genetic disorders, which can be chronic, debilitating, and life-threatening conditions. Lipodystrophy is one example of this ^4^, and is characterised by altered adipose tissue mass, distribution or function^5^. Individuals with severe forms of lipodystrophy, such as congenital generalised lipodystrophy type 2 (CGL2), have significantly reduced adipose mass and develop complex metabolic complications, including insulin resistance, lipoatrophic diabetes, hypertriglyceridemia, and hepatic steatosis ^6^. Lipodystrophy also substantially impacts quality of life for individuals with this disorder and has been shown to reduce lifespan by more than thirty years ^7^. Due to its scarcity, a lack of understanding and insufficient scientific research into this disorder, effective therapeutic treatment options for lipodystrophy remain very limited ^8^.

Adipose tissue has recently emerged as an attractive target for AAV vectors. Intensive research efforts have identified specific AAV serotypes and novel recombinant AAV vectors that can effectively target adipose tissue depots ^9^. These studies have also provided proof of concept that AAV-mediated gene therapy could effectively be utilised to correct disorders of adipocyte dysfunction, such as leptin deficiency ^10, 11^, and reverse metabolic complications in a preclinical model of a known human disorder. We recently revealed that systemic AAV-mediated gene therapy, restoring functional expression of the defective *Bscl2* gene, is also capable of restoring adipose tissue development and metabolic health in a preclinical mouse model of CGL2 ^12^. This approach utilised AAV8 vectors to drive transgene expression using the strong ubiquitous cytomegalovirus (CMV) promoter. However, clinically, tissue-selective promoters are now favoured for gene therapy when delivered systemically ^2, 3^. Whilst substantial advances have been made in gene therapies targeting other tissues, research exploring AAV-mediated gene therapy for adipose tissue disorders, and in particular lipodystrophy, remains extremely limited ^9^.

In this study we used the mini/aP2 promoter to examine whether adipose tissue-selective AAV gene therapy is capable of rescuing adipose development and restoring metabolic health in a preclinical mouse model of CGL2. We also evaluated the effects of liver-selective expression using the thyroxine binding globulin (TBG) promoter ^13, 14^. This revealed that adipose, but not liver-selective re-expression of functional human *BSCL2*, is highly effective in restoring adipose development and metabolic health, thereby offering an effective and refined therapeutic intervention strategy for lipodystrophies.

## MATERIALS AND METHODS

### Ethics approval statement

Procedures on wild type (WT) and seipin knockout (SKO) mice were approved by the University of Aberdeen Ethics Review Board and performed under the Animals (Scientific Procedures) Act 1986 under project license (PPL: P1ECEB2B6) approved by the UK Home Office.

### Animal studies

SKO mice were generated as previously described and maintained on a C57BL/6J background ^15^. Power calculations were used to estimate sample size based on anticipated changes in blood glucose levels from values and standard deviations obtained during previous procedures. In order to attain statistical significance of p <0.05 with a power level of 80%, a sample size of eight mice per group was anticipated. Mice had *ad libitum* access to water and chow diet (CRM (P) 801722, Special Diets Services) unless otherwise stated. Mice were group-housed in home cages at 20-24°C, 45-65% humidity and exposed to a 12hr/12hr light-dark period. Body composition was measured using the EchoMRI™-500 analyser (Zinsser Analytic GmbH). Blood glucose levels were determined by glucometer readings (AlphaTrak^®^ II, Zoetisus) from tail punctures. Tissues were rapidly dissected post-mortem, frozen in liquid nitrogen and stored at −70°C.

### AAV administration

Prior to AAV administration, 15- to 17-week-old male and female SKO mice were randomised then intraperitoneally (I.P.) injected with 1×10^12^ genome copies of AAV8 expressing the long form of human *BSCL2* (NM_001122955.4) under the cytomegalovirus (CMV), mini/aP2 (aP2) or thyroxine binding globulin (TBG) promoter (AAV8-CMV-hBSCL2 (VB191203-2053snp), AAV8-mini-aP2-hBSCL2-miR-122 (VB211102-1003sae), AAV8-TBG-hBSCL2 (VB220221-1474ujt), VectorBuilder) or AAV8-CMV-eGFP (VB010000-9394npt, VectorBuilder) vectors. For low titer studies, 9- to 12-week-old male and female SKO mice were I.P. injected with 1×10^10^ genome copies of AAV8-mini-aP2-hBSCL2-miR-122 or AAV-TBG-hBSCL2 vectors. Investigators were not blinded to the group allocation during experiments and/or when assessing the outcomes. The I.P. route of administration can occasionally be unreliable, for example inadvertent injection into the gut. Mice determined to have not responded to AAV treatment were therefore excluded from all analysis.

### Tolerance tests

Oral glucose and insulin tolerance tests were performed five and eight weeks after AAV administration respectively. Male and female mice were placed in clean cages and food was withheld for 5 hours. Basal glucose levels (0 min) were determined by glucometer readings (AlphaTrak® II, Zoetisus) from tail punctures. Mice were then administered either 2 g/kg d-glucose (Sigma-Aldrich) bolus by gavage or 0.75 IU/kg human insulin (ActRapid, NovoNordisk) by I.P. injection. Blood glucose levels were monitored at 15-, 30-, 60- and 90-min. Mice had *ad lib* access to water throughout.

### Immunohistochemistry

Liver and adipose tissues were fixed in 10% formalin, embedded in paraffin and 5 µm sections cut. Sections were deparaffinised in xylene, and rehydrated. Epitope retrieval was performed using 10 mM Sodium Citrate Buffer (pH 6, 0.05% Tween) and incubating at 126°C for 3 min. For endogenous peroxidase quenching, sections were treated with 3% H_2_O_2_ in methanol for 15 minutes followed by 2X H2O, 2X TBS-Triton (0.025% Triton) and 2X TBS-Tween-20 (0.1% Tween-20) 5-minute washes each. After blocking in 3% BSA in TBS-T for 1 hour at room temperature, sections were incubated overnight with primary antibody at 1:500 (BSCL2/Seipin, #23846, Cell Signaling) in 1% BSA in TBS-T at 4°C. Sections were washed 3X 5 min with TBS-T and incubated with secondary antibody at 1:500 (#7074, Cell Signaling) in 1% BSA in TBS-T for 1 hour at room temperature. After 3 washes with TBS-T for 5 min, DAB staining was performed as per manufacturer’s protocol (Vector, DAB Peroxidase Substrate, Cat: SK-4100). Slides were dehydrated and coverslips were mounted with CV Mount (Leica, 14046430011).

### H&E staining

Deparaffinised 5 µm liver and adipose tissue sections fixed in 10% formalin were rinsed in H2O and stained with hematoxylin solution (GHS316-500ml, Sigma-Aldrich) for 5 minutes and washed with H_2_O for 2 minutes. Destaining/Differentiation was performed with 1% acid alcohol for 30 seconds and 1 minute H_2_O, followed by bluing with saturated lithium carbonate solution for 1 minute. Sections were washed with H_2_O and stained with eosin (1% eosin, 2% phloxine, 2.5% CaCl_2_) for 2 minutes. Slides were dehydrated by washing in H_2_O for 20 seconds, 2X 70% ethanol, 2X 100% Ethanol and 2X 2 minutes incubation in xylene. Coverslips were mounted with CV Mount (Leica, 14046430011).

### Serum analysis

Blood was collected in SST™ amber tubes (BD Microtainer^®^) from five-hour fasted mice by cardiac puncture, inverted and incubated at room temperature for 30 minutes. Samples were centrifuged at 12,000 × g for 10 minutes and serum collected. Insulin, glucose, adiponectin, leptin, Fgf21, alanine transaminase (ALT) and aspartate transaminase (AST) analysis was performed at the Core Biochemical Assay Laboratory (Cambridge, UK). Serum triglyceride (TG) levels were determined using the Triglyceride Liquid Assay (Sentinel Diagnostics). Quantitative insulin sensitivity check index (QUICKI) was calculated as previously described ^16^. QUICKI = 1/[log(I_0_) + log(G_0_)], where I_0_ is fasting insulin (µU/mL) and G_0_ is fasting glucose (mg/dL).

### Gene expression

RNA was extracted from frozen tissues using the RNeasy mini kit (Qiagen). Equal quantities of total RNA were DNase I treated (Sigma), reverse transcribed using the Tetro cDNA Synthesis Kit following the manufacturer’s protocol (Meridian Biosciences). Real-time quantitative PCR was performed on the CFX384 Touch™ Real-Time PCR Detection System (BioRad). No template and no reverse transcriptase controls were performed for every gene analysed. The geometric mean of three stable reference genes (*Nono*, *Ywhaz* and *Hprt*) was used for normalisation.

### Liver triglyceride assay

Frozen liver tissue samples were weighed, homogenised in 1 ml of PBS, and kept on ice. Liver lysates were centrifuged at 6,000 × g for 10 minutes at 4°C. Supernatants were collected and TG levels determined using the Triglyceride Liquid Assay (Sentinel Diagnostics) and normalised to tissue weight.

### Western blot

Frozen tissues were homogenised in RIPA buffer containing cOmplete protease inhibitor (Roche). Protein concentrations were determined by BCA assay (Thermo). SDS-PAGE was performed using equal quantities of protein and transferred to PVDF membrane. All biological replicates have been assessed for transgene expression. For tissue panel analysis, human BSCL2 protein levels were examined in multiple tissues from a single representative mouse for each treatment. Antibodies used at a dilution of 1 in 1-2000 included anti-BSCL2/Seipin, which is only capable of reacting with human BSCL2/Seipin (#23846, Cell Signaling) and anti-Calnexin (ab75801, Abcam). Anti-Rabbit HRP secondary antibody was used at a dilution of 1 in 5000 (#7074, Cell Signaling) and visualised using enhanced chemiluminescence (Immobilon Crescendo Western HRP Substrate, Millipore, #WBLUR0500).

### Statistical analyses

All data are presented as mean ± SD. Biological data was assessed for normality using the Shapiro-Wilk normality test. Experiments where individual biological data was determined to have excessively deviated from a normal distribution was assessed for lognormal distribution and then the entire dataset was transformed to logarithms prior to analysis. Data was analysed by one-way ANOVA with Tukey post-hoc test or two-way repeated measures ANOVA with Bonferroni post-hoc test as appropriate using GraphPad Prism. If SDs were significantly different as determined by the Brown-Forsythe test, Brown-Forsythe and Welch ANOVA tests were performed. A *P*-value <0.05 was considered as statistically significant.

## RESULTS

### Gene therapy mediated reversal of hyperglycaemia in SKO mice

We previously revealed that gene therapy effectively restores adipose tissue development and metabolic health in a preclinical mouse model of CGL2 ^12^. This approach utilised AAV8 vectors containing the human *BSCL2* gene driven by the strong ubiquitous CMV promoter (AAV8-CMV-hBSCL2). Use of the human *BSCL2* gene was employed to allow for specific detection of our transgene in target tissues. To examine if tissue-selective gene therapies could be effective, we designed AAV8 vectors containing the mini/aP2 ^13^ or TBG ^14^ promoters (Figure 1A) to drive human *BSCL2* gene expression selectively in adipose tissue or liver, respectively. The mini/aP2 vector also contained four copies of the liver-specific microRNA-122 (miR-122) target sequence, to restrict off-target hepatic transgene expression.

**Figure 1:**
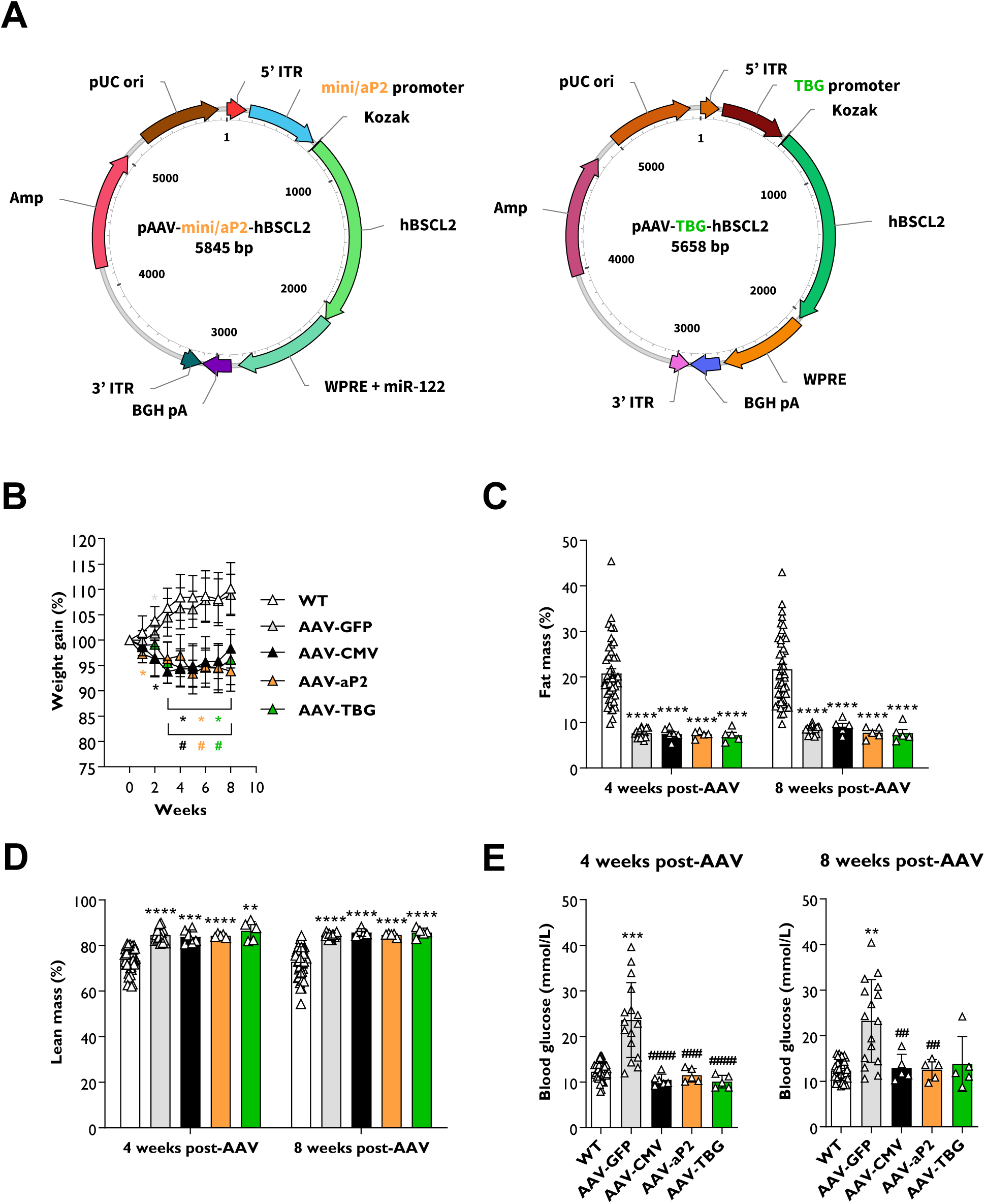
Tissue-selective gene therapies reverse hyperglycaemia in SKO mice. (**A**) Viral vector plasmid design to overexpress the long form of the human *BSCL2* transcript (NM_001122955.4) driven by the adipose-selective mini/aP2 (pAAV8-mini/aP2-hBSCL2) or liver-selective thyroxine binding globulin (pAAV8-TBG-hBSCL2) promoters. (**B**) Weight gain progression in wild type (WT) and seipin knockout (SKO) mice after I.P. injection of 1×10^12^ genome copies of AAV8 vectors containing GFP (AAV-GFP), or human BSCL2 under control of the CMV (AAV-CMV), mini/aP2 (AAV-aP2) or TBG promoter (AAV-TBG). Whole body fat mass (**C**) and whole body lean mass (**D**) levels normalised to body weight and assessed by EchoMRI four and eight weeks after AAV administration. (**E**) Blood glucose levels in *ad lib* fed WT, AAV-GFP, AAV-CMV, AAV-aP2 and AAV-TBG treated mice at four and eight weeks after administration of gene therapy. All data are biological replicates presented as the mean ± SD, n = 44 (WT), 14 (AAV-GFP), 5-6 (AAV-CMV), 5 (AAV-aP2) and 5 (AAV-TBG) mice per group, * p<0.05, ** p<0.01, *** p<0.001 and **** p<0.0001 vs WT, #p<0.05, ## p<0.01, ### p<0.001 and #### p<0.0001 vs AAV-GFP.

Cohorts of 15- to 17-week-old male and female seipin knockout (SKO) mice were randomised and I.P. injected with 1×10^12^ genome copies of AAV8-CMV-hBSCL2 (AAV-CMV), AAV8-mini/aP2-hBSCL2 (AAV-aP2), AAV8-TBG-hBSCL2 (AAV-TBG) or AAV8-CMV-eGFP (AAV-GFP) as a control. Mice were fed a chow diet for 8 weeks and physiological and metabolic measurements were performed (Figure S1A). Similar to our previous observations ^12^, weight gain in wild type (WT) and AAV-GFP injected mice was comparable, and AAV-CMV injected mice failed to gain weight after AAV administration (Figure 1B). Interestingly, weight gain also ceased in both AAV-aP2 and AAV-TBG treated mice, comparable to AAV-CMV treated mice (Figure 1B). EchoMRI analysis revealed significantly decreased body fat mass (Figure 1C) and increased lean mass (Figure 1D) in SKO mice when normalised to body weight compared with WT mice. This was observed both at 4- and 8-weeks after AAV injection, with no significant differences between treatment groups. As expected, examination of *ad lib* blood glucose levels revealed AAV-GFP injected mice were hyperglycaemic, and this was significantly reversed by AAV-CMV injection at 4- and 8-weeks (Figure 1E). Similar significant reductions in blood glucose levels were also apparent in AAV-aP2 and AAV-TBG treated SKO mice 4-weeks post-AAV and in AAV-aP2 treated mice 8-weeks post AAV, with no significant differences observed when compared with WT mice (Figures 1E).

These findings indicate that gene therapy designed to selectively target adipose tissue appears as effective as untargeted CMV driven gene therapy at reversing hyperglycaemia in a preclinical mouse model of CGL2. Surprisingly however, gene therapy using the liver-selective TBG promoter also gave similar results.

### Effect of gene therapies on metabolic complications in SKO mice

To determine if tissue-selective gene therapies had additional beneficial effects, we examined metabolic complications present in SKO mice. Oral glucose and insulin tolerance tests were significantly impaired in AAV-GFP treated mice compared to WT controls (Figure 2A+B). This was completely reversed in mice receiving either AAV-aP2 or AAV-TBG vectors (Figure 2A+B). Fasted circulating insulin levels were significantly elevated in AAV-GFP treated mice compared with WT controls (Figure 2C). Both mini/aP2 and TBG promoter driven expression of hBSCL2 restored fasting insulin to similar levels observed in WT controls (Figure 2C). Fasting glucose levels were not significantly different between any of the groups examined (Figure 2D). Quantitative insulin-sensitivity check index (QUICKI) analysis was significantly decreased in AAV-GFP treated mice compared with WT mice, confirming insulin resistance (Figure 2E). Again, both AAV-aP2 or AAV-TBG gene therapies completely restored insulin sensitivity by this measure in SKO mice (Figure 2E).

**Figure 2:**
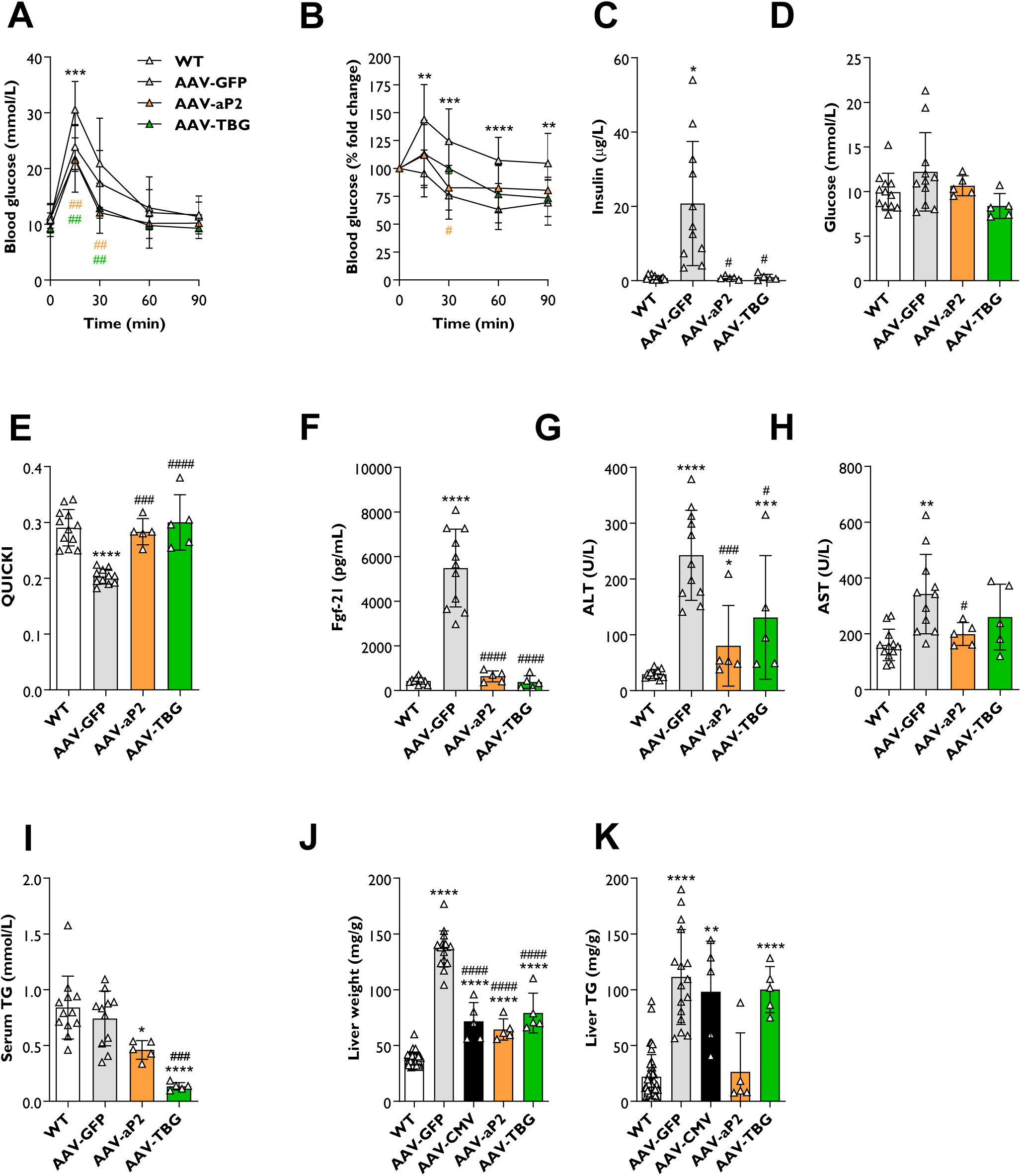
Tissue-selective gene therapies improve metabolic complications in SKO mice. Oral glucose (**A**) and insulin (**B**) tolerance tests in WT, AAV-GFP, AAV-aP2 and AAV-TBG treated mice. The glucose bolus (2 g/kg) was given by gavage. Human insulin (0.75 IU/kg) was administered by I.P. injection. Serum insulin (**C**), glucose (**D**), quantitative insulin sensitivity check index (QUICKI) analysis (**E**), Fgf21 (**F**), ALT (**G**), AST (**H**) and triglyceride (TG) levels (**I**) in WT, AAV-GFP, AAV-aP2 and AAV-TBG treated mice fasted for five hours and eight weeks after AAV administration. Liver weights normalised to body weight (**J**) and liver TG levels (**K**) in WT, AAV-GFP, AAV-aP2 and AAV-TBG treated mice eight weeks after AAV administration. All data are biological replicates presented as the mean ± SD, n = 32 (WT), 11 (AAV-GFP), 5 (AAV-aP2) and 5 (AAV-TBG) mice per group, for A-I, n = 44 (WT), 16 (AAV-GFP), 5 (AAV-CMV), 5 (AAV-aP2) and 5 (AAV-TBG) mice per group for J-K, *p<0.05, ** p<0.01, *** p<0.001 and **** p<0.0001 vs WT and # p<0.05, ## p<0.01, ### p<0.001 and #### p<0.0001 vs AAV-GFP.

We next examined key circulating serum factors. AAV-GFP treated mice had significantly elevated circulating Fgf21 levels compared with WT mice (Figure 2F). This was completely normalised in mice receiving AAV-aP2 and AAV-TBG vectors (Figure 2F). Markers of liver damage, alanine aminotransferase (ALT) and aspartate transaminase (AST), were significantly elevated in AAV-GFP treated mice (Figure 2G+H). Both AAV-aP2 and AAV-TBG gene therapies significantly reduced circulating ALT, however remained significantly elevated compared to WT levels (Figure 2G). Both treatments also reduced circulating AST levels, which was statistically significant in AAV-aP2 treated mice (Figure 2H). Serum triglyceride (TG) levels were not significantly different between WT and AAV-GFP treated mice (Figure 2I). However, a significant reduction of serum TG was evident in AAV-aP2 treated mice, with an even greater decrease observed in SKO mice treated with AAV-TBG vectors (Figure 2I).

We also examined the effect of gene therapies on liver weights, liver TG levels and hepatic gene expression markers. Significantly increased liver weights and liver TG levels were observed in AAV-GFP treated mice compared to WT controls (Figure 2J+K). Liver weights were significantly decreased in SKO mice treated with AAV-aP2 and AAV-TBG, similar to the effect observed in SKO mice treated with AAV-CMV (Figure 2J). These significant reductions in liver weight predominantly account for the reduced weight gain observed in AAV treated mice (Figure 1B). Despite reductions in liver weight, no significant differences were observed in liver TG levels measured per gram of tissue between AAV-GFP, AAV-CMV or AAV-TBG treated mice (Figure 2K). Curiously however, liver TG levels per gram of tissue were reduced in AAV-aP2 injected mice and normalised to WT levels, although this did not reach statistical significance (Figure 2K). Expression levels of genes known to be dysregulated in the livers of SKO mice such as *Scd1*, *Fgf21*, *Fabp4* and *Pparg* were significantly upregulated in AAV-GFP treated SKO mice compared with WT controls (Figure S1B+C). These increases were reversed by either AAV-aP2 or AAV-TBG injection to SKO mice, apart from for *Scd1* which remained elevated in mice injected with the adipose tissue-selective AAV-aP2 vector. Expression of *Ppara* was not significantly altered in SKO mice or either of the treatment conditions (Figure S1B+C).

Overall, these findings reveal that adipose tissue-selective AAV-aP2 gene therapy can significantly improve multiple metabolic complications in SKO mice. Surprisingly, AAV-TBG, which is intended to drive liver-selective expression of human *BSCL2*, produced similar effects when delivered at the same titer.

### Evaluation of AAV vector tissue specificity in SKO mice

Previous reports indicate that hepatic *Bscl2* deficiency does not play a significant role in the development of metabolic disease in SKO mice ^14, 17^. It is therefore unclear why AAV-TBG treatment restored metabolic health in SKO mice if this provided liver-selective restoration of human BSCL2 as intended. We therefore characterised in detail the tissue specificity of our gene therapy vectors.

Human *BSCL2* mRNA was detectable in the livers of all SKO mice receiving AAVs encoding hBSCL2. However, AAV-aP2 injected mice expressed only modest levels of hBSCL2 compared to AAV-CMV treated SKO mice (Figure 3A). In contrast, AAV-TBG injection led to dramatically higher hBSCL2 mRNA expression in the livers of SKO mice, more than thirty-fold higher than in AAV-CMV treated mice. Histological staining for human BSCL2 protein confirmed the robust expression in AAV-TBG treated mice, with more modest expression in AAV-CMV treated mice (Figure 3B). Despite detecting human *BSCL2* mRNA in AAV-aP2 treated samples, albeit at low levels (Figure 3A), human BSCL2 protein was not detectable in liver sections from these mice (Figure 3B). This is likely due to the incorporation of the liver-specific miR-122 target sequence in this vector (Figure 1A).

**Figure 3:**
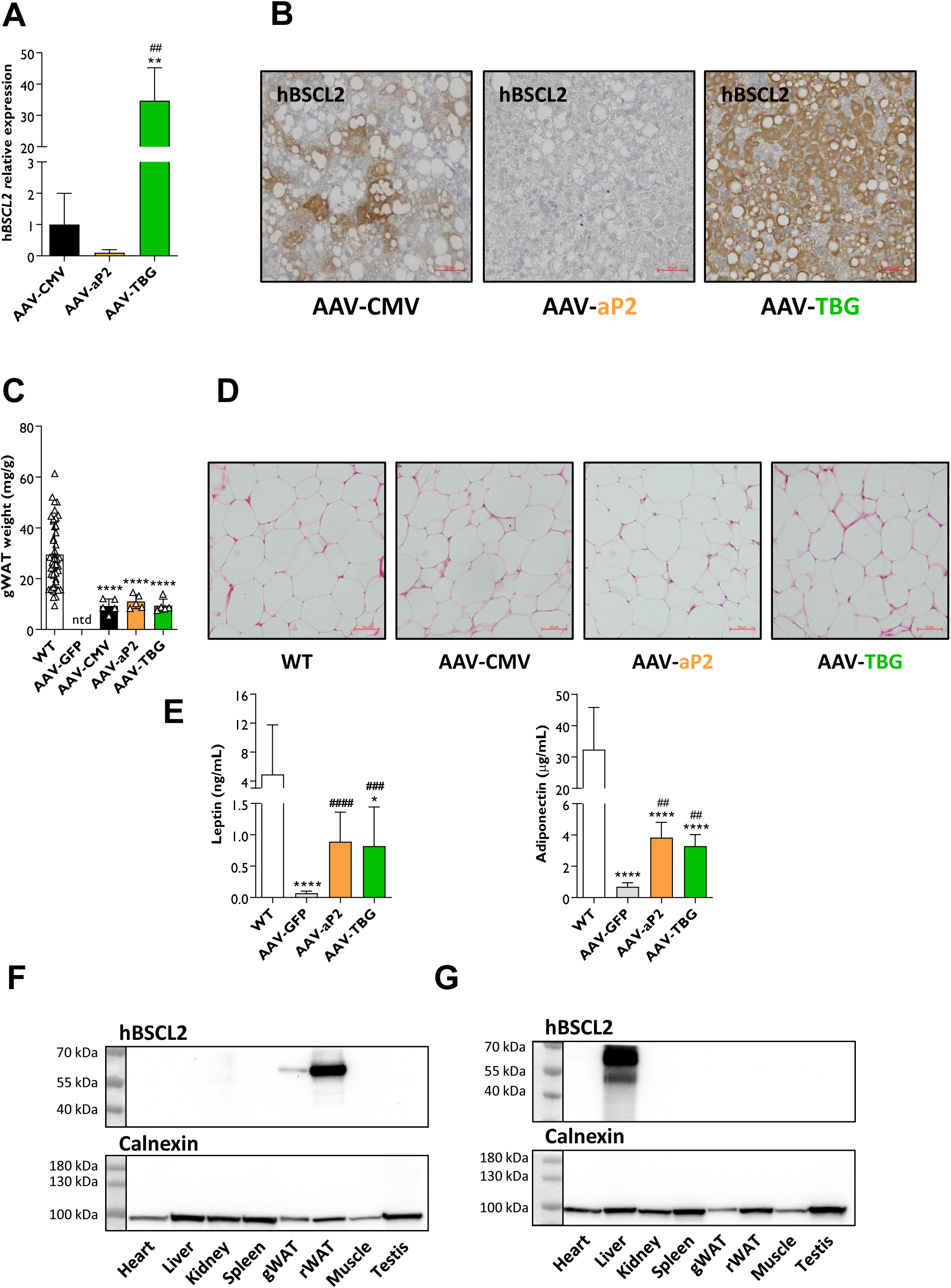
Evaluation of AAV vector tissue specificity in SKO mice. Relative human *BSCL2* gene expression levels (**A**) and representative antibody staining of human BSCL2 protein (**B**) from livers of AAV-CMV, AAV-aP2 and AAV-TBG treated mice eight weeks after AAV administration. Gonadal white adipose tissue (gWAT) weights (**C**) and representative H&E sections of gWAT (**D**) from WT, AAV-CMV, AAV-aP2 and AAV-TBG treated mice. (**E**) Circulating serum levels of leptin and adiponectin in WT, AAV-GFP, AAV-aP2 and AAV-TBG injected mice fasted for five hours. Western blot tissue panel analysis of human BSCL2 protein levels in AAV-aP2 (**F**) and AAV-TBG (**G**) injected mice. Data in A, C and E are biological replicates presented as the mean ± SD, n = 5 (AAV-CMV), 5 (AAV-aP2) and 5 (AAV-TBG) mice per group for A, n = 44 (WT), 5 (AAV-CMV), 5 (AAV-aP2) and 5 (AAV-TBG) mice per group for C, n = 12 (WT), 10-11 (AAV-GFP), 5 (AAV-aP2) and 5 (AAV-TBG) for E. * p<0.05, ** p<0.01, and **** p<0.0001 vs WT, ## p<0.01, ### p<0.001 and #### p<0.0001 vs AAV-GFP by one-way ANOVA, ntd = no tissue dissected. Scale bars represents 50 µm.

As reported previously ^12^, visceral adipose tissue depots were completely absent in AAV-GFP injected mice, but substantially restored in AAV-CMV injected mice (Figure 3C+S1D). Similar quantities of gonadal and retroperitoneal white adipose tissue (gWAT and rWAT respectively) were dissectible from AAV-aP2 treated mice. Surprisingly, this was also the case for AAV-TBG injected mice (Figure 3C+S1D). H&E sections from each AAV treatment revealed that gWAT adipocyte size and cell morphology were similar to that observed in WT mice (Figure 3D). We confirmed the absence of murine *Bscl2* mRNA and observed substantial expression of key murine adipocyte markers (*Pparg*, *Leptin* and *Adiponectin*) in restored gWAT of AAV-aP2 and AAV-TBG injected mice (Figure S1E+F). Consistent with the rescue of functional adipose tissue, circulating adipose derived hormones leptin and adiponectin were found to be significantly increased in both AAV-aP2 and AAV-TBG injected SKO mice compared to AAV-GFP injected controls (Figure 3E).

Interestingly, human *BSCL2* mRNA expression was evident in gWAT of both AAV-aP2 and AAV-TBG injected mice (Figure S1E+F) indicating that the latter can drive adipose expression at this titer despite using the liver-selective TBG promoter. To examine the tissue specificity of each vector further, we assessed human BSCL2 protein levels in multiple tissues from AAV-aP2 and AAV-TBG treated mice by western blot analysis. BSCL2 expression was present in gWAT and more highly expressed in rWAT of AAV-aP2 injected mice, but not detected in the liver or other tissues examined (Figure 3F). In contrast, BSCL2 was highly expressed in the liver, and appeared undetectable in gWAT and rWAT of AAV-TBG injected mice (Figure 3G). The high level of BSCL2 expression in liver of AAV-TBG injected mice may have masked low level expression in other tissue. We therefore repeated western blot analysis excluding the liver and indeed observed low level off-target expression of human BSCL2 in gWAT and rWAT, along with muscle and testis (Figure S1G). Finally, we compared BSCL2 expression in gWAT from AAV-CMV, AAV-aP2 and AAV-TBG treated SKO mice. Encouragingly, the use of the adipose tissue-selective promoter (AAV-aP2) resulted in similar human BSCL2 protein expression to that found when using the strong ubiquitous CMV promoter (AAV-CMV). In contrast, little to no hBSCL2 protein was detected in the AAV-TBG treated mice when directly compared (Figure S1H).

Together, these findings indicate that high adipose tissue specificity appears to have been achieved for the AAV-aP2 vector. However, low level off-target transgene expression in other tissues including adipocytes or their precursors was present when using the AAV-TBG vector administered at high titers (1×10^12^ genome copies).

### Examining tissue-selective gene therapies at low AAV titers in SKO mice

Our findings confirm previous reports that adipose tissue restoration is responsible for improvements to metabolic health in SKO mouse models ^18–20^. However, it remains unclear if liver-selective gene therapy could be of therapeutic benefit. We previously reported that AAV8 vectors target the liver, but not adipose tissue depots when I.P. injected at 1×10^10^ genome copies ^12^. Therefore, we utilised this lower dose to determine whether liver-selective gene therapy was indeed effective.

Cohorts of 9- to 12-week-old male and female SKO mice were I.P. injected with 1×10^10^ genome copies of AAV-aP2 or AAV-TBG vectors. Mice were then fed a chow diet for 4 weeks and physiological and metabolic measurements were performed. At this lower dose, both AAV-aP2 and AAV-TBG treated mice gained weight that was comparable and not significantly different to WT controls (Figure 4A). Examination of blood glucose levels in mice fasted for 5 hours and re-fed chow for 1 hour revealed that AAV-aP2 and AAV-TBG treated mice remained hyperglycaemic at 2- and 4-weeks post-injection (Figure 4B). Postmortem serum TG levels were also not significantly different between WT and AAV injected mice (Figure 4C). Upon dissection, gWAT (Figure 4D) and rWAT (images not shown) was not evident in either AAV-aP2 or AAV-TBG treated mice. Despite confirming the presence of human BSCL2 protein in livers of AAV-TBG treated mice by western blot analysis (Figure 4E), no significant improvements to liver weights (Figure 4F) or liver TG levels (Figure 4G) were apparent in either group. Similarly, no significant improvements were observed to the dysregulated liver markers *Scd1* or *Pparg*, which remained elevated under these conditions (Figure 4H).

**Figure 4:**
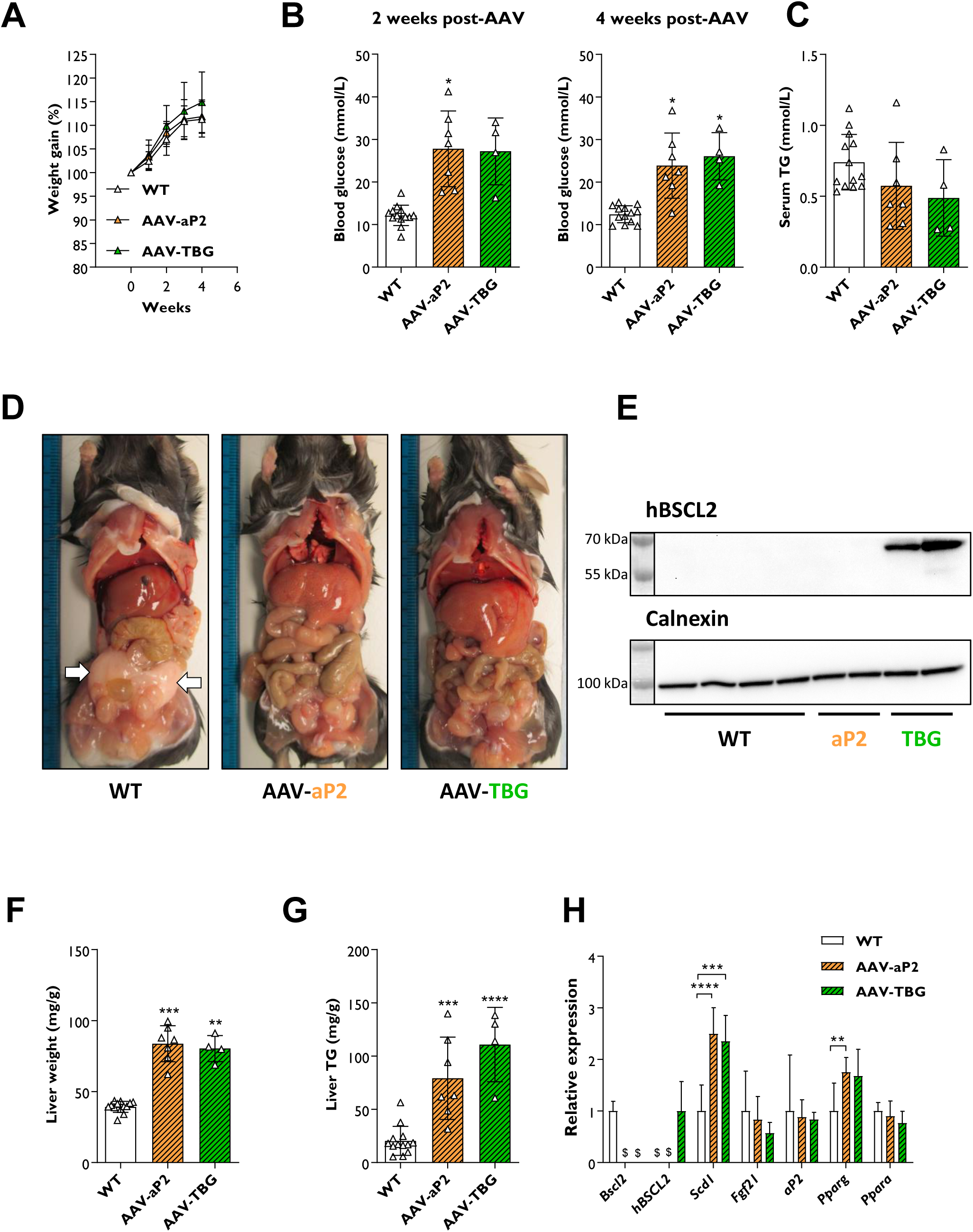
Examining tissue-selective gene therapies at low AAV titers in SKO mice. (**A**) Weight gain progression in WT and SKO mice after I.P. injection of 1×10^10^ genome copies of AAV-aP2 or AAV-TBG. (**B**) Blood glucose levels in WT, AAV-aP2 and AAV-TBG treated mice fasted for five hours and re-fed chow for one hour at two and four weeks after AAV administration. Serum TG levels (**C**), postmortem dissection images (**D**) and western blot analysis of human BSCL2 protein levels in liver (**E**) of WT, AAV-aP2 and AAV-TBG treated mice four weeks after AAV administration. White arrows in D highlight gWAT depot in WT mice. Liver weights normalised to body weight (**F**), liver TG levels (**G**) and relative gene expression levels of metabolic markers in the liver (**H**) of WT, AAV-aP2 and AAV-TBG treated mice four weeks after AAV administration. All data are biological replicates presented as the mean ± SD, n = 13 (WT), 7 (AAV-aP2) and 4 (AAV-TBG) mice per group, * p<0.05, ** p<0.01, *** p<0.001 and **** p<0.0001 vs WT, $ = not detected.

Overall, these findings reveal that the liver-selective AAV-TBG gene therapy vector when administered at 1×10^10^ genome copies is unable to improve metabolic dysfunction that is present in a preclinical mouse model of CGL2.

## DISCUSSION

Globally, rare disorders are thought to impact around 6% of the population ^21^. However, the scarcity of individual conditions poses significant challenges to the development and implementation of viable and effective therapeutic solutions. Gene therapy has emerged as a practical and transformative medical intervention aimed at preventing, treating, and potentially curing an array of genetic and acquired disorders ^1–3^. Its efficacy is particularly pronounced in addressing rare monogenic disorders, which can be chronic, debilitating, and life-threatening conditions.

Lipodystrophy is a rare disorder ^4^ characterised by abnormal or deficient adipose tissue formation and distribution in the body ^5^. The only current dedicated treatment strategy for generalised lipodystrophy is metreleptin ^22^ which can be highly effective, but it is expensive and not always widely available. Other pharmacological therapies are currently being investigated and there is evidence that GLP-1R agonists could be useful in the treatment of CGL2 ^23^. Due to the severity of this disorder, alternative and invasive strategies have also been explored for generalised and partial lipodystrophies, including bariatric surgery ^5^ and a recent first-in-literature report of a liver transplantation ^24^. However, current therapies only manage the severe metabolic complications that develop, and none can restore adipose tissue development or functionality ^8^.

Several studies have revealed that adipose tissue can be effectively targeted by multiple AAV serotypes ^9^. Despite the complete absence of adipose tissue in mice with severe lipodystrophy, we recently revealed that AAV vectors can restore adipose tissue development and metabolic health in a preclinical mouse model of CGL2 ^12^. This approach utilised AAV8 vectors to drive transgene expression using the strong ubiquitous CMV promoter, which targeted multiple tissues including the liver and rescued adipose tissue. Previous research indicates that restoration of adipose tissue alone is responsible for the improvements to glucose homeostasis in SKO mice ^18–20^. However, a recent study highlighted that AAV-mediated overexpression of *Bscl2* in the liver alleviated high-fat diet-induced hepatic steatosis in WT mice ^25^, potentially indicating beneficial liver-mediated effects. Therefore, to investigate the capabilities of gene therapy for lipodystrophy further, we evaluated the efficacy of both adipose and liver tissue-selective gene therapies using the mini/aP2 ^13^ and TBG ^14^ promoters respectively.

It has been reported that adipose-selective mini-promoters are significantly less potent than ubiquitous promoters ^9, 13^. Despite this, AAV vectors harbouring the mini/aP2 promoter effectively restored visceral adipose tissue development and metabolic health in SKO mice, when used at the same dose (1×10^12^) as AAV vectors containing the CMV promoter. Transgene expression appeared highly selective for adipose tissue, and off-target hepatic transgene expression was prevented through the incorporation of the liver-specific microRNA target sequence miR-122 into the AAV vector ^26^. Therefore, our findings indicate that adipose tissue-selective AAV-mediated gene therapy for lipodystrophy is feasible and could provide an effective and refined form of therapeutic intervention without the need to use strong ubiquitous promoters. This is particularly encouraging as recent trends in AAV clinical trials indicate an increased usage of tissue-selective promoters, particularly when administration is by systemic delivery ^2, 3^.

Curiously, AAV vectors containing the liver-selective TBG promoter were equally effective at restoring metabolic health in SKO mice when administered at dose of 1×10^12^ genome copies. If liver-mediated gene therapy could restore adipose tissue development and metabolic health in lipodystrophy, this would be highly attractive due to the extensive prior knowledge and application of this approach that has been gained from numerous clinical trials targeting this organ ^2^. While there is evidence that overexpressing *Bscl2* in the liver may be beneficial in high-fat diet fed mice ^25^, the literature indicates that hepatic *Bscl2* deficiency does not play a significant role in the development of metabolic disease in SKO mice ^14, 17^. Unexpectedly, when using the TBG promoter, visceral adipose tissue depots developed to a similar extent seen when using the CMV and mini/aP2 promoters. This is despite the fact that transgene expression appeared to be highly liver-selective by western blot analysis.

One explanation for the effects observed when using AAV8-TBG vectors at high doses is a transient/low level of *BSCL2* transgene expression present in adipose progenitor cells, enabling adipose tissue development to occur. Indeed, adipose-specific ablation of *Bscl2* in adult mice does not result in the severe lipodystrophy observed in SKO mice ^27–29^. Therefore, adipose tissue depots can be maintained when Bscl2 protein levels are low or absent. Confirming this possibility, a recent study identified that standard screening methods underreport AAV8-mediated transduction when examining gene editing after AAV-CRISPR delivery ^30^. The authors revealed that AAV8 vectors can target numerous and previously unknown cell and tissue types through transient/low level transgene expression, which have previously been missed when examining transduction patterns based on high and stable expression of reporter genes. In agreement with this, we were able to detect human *BSCL2* mRNA in gWAT and low human BSCL2 protein expression in gWAT, rWAT, muscle and testis in SKO mice injected with AAV-TBG. We previously reported that AAV8 transgene expression was detectable in the liver, but not adipose tissue when administered at a dose of 1×10^10^ genome copies ^12^. Using this lower AAV titer, we confirmed the presence of BSCL2 protein in the liver, however adipose tissue development and improved metabolic health was no longer observed in SKO mice. Therefore, liver-mediated gene therapy for CGL2 is unlikely to be a feasible approach to improve metabolic health.

Our data have revealed novel insights into the feasibility of tissue-selective gene therapy for lipodystrophy disorders. However, several issues and challenges remain if this is to become a realistic mode of therapeutic intervention. Whilst the use of an adipose tissue-selective promoter AAV vector was effective, the amount of visceral adipose tissue development was no greater than that observed with a strong ubiquitous promoter. Additionally, this approach again failed to stimulate subcutaneous adipose tissue development, as observed in our previous studies ^12^. Far more research will be necessary to determine the optimal requirements to target and stimulate adipose tissue development in this depot. This will include examining different routes of administration such as direct injection, intravenous delivery, and oral administration ^9, 31^, which we have not yet performed through detailed studies in our preclinical mouse model of lipodystrophy. The development and assessment of novel AAV serotypes ^32^ and exploration of alternative adipose tissue/cell-specific promoters ^10^ will also be critical. Furthermore, despite an extensive history of AAV usage in the clinical setting, to our knowledge, no clinical trials to date have aimed to directly target adipose tissue ^2, 3^. Therefore, no clinical information is available to suggest that what has been observed in mouse models ^9, 12^ could be effectively translated to human disorders of adipose tissue dysfunction.

Overall, our findings have evaluated the potential of tissue-selective AAV-mediated gene therapy for lipodystrophy disorders. Our data conclusively prove that liver-selective AAV-mediated gene therapy is unfortunately not an effective strategy for CGL2 and should not be considered further as a therapeutic intervention. However, we reveal that usage of an adipose tissue-selective mini-promoter within AAV vectors is an effective form of therapeutic intervention to restore metabolic health in severe forms of lipodystrophy such as CGL2. Continued efforts should be made to develop this form of therapy to provide an effective treatment for this rare and devastating disorder.

## DATA AVAILABILITY

The datasets generated and/or analysed during the current study are available from the corresponding author on reasonable request.

## ACKNOWLEDGEMENTS

The authors would like to thank the staff at the University of Aberdeen’s Microscopy and Histology Core Facility and the Medical Research Facility for support with animal breeding and maintenance. The manuscript figure was created with BioRender.com. This research was supported by funding from Diabetes UK RD Lawrence Fellowship (21/0006280) awarded to G.D.M, from the Biotechnology and Biological Sciences Research Council (BB/V015869/1) to JJR and a University of Aberdeen Doctoral Training Grant to M.T. NS is supported by a BBSRC East of Scotland Bioscience Doctoral Training Partnership (EASTBIO) PhD studentship.

## AUTHOR CONTRIBUTIONS

M.T. acquired data and interpreted results, A.R. and N.S. acquired data, W.H., M.D. and J.J.R. assisted with design of the work, G.D.M. conceived the study, designed, acquired, and interpreted the results and drafted the manuscript. All authors have revised and approved the final version of the manuscript and agree to be accountable for all aspects of the work. G.D.M is the guarantor of this work and, as such, had full access to all the data in the study and takes responsibility for the integrity of the data and the accuracy of the data analysis.

