## Supplemental Figure 1 for "Preclinical evaluation of tissue-selective gene therapies for congenital generalised lipodystrophy"

**Figure S1:** Tissue-selective gene therapy in lipodystrophy. (**A**) Graphical illustration of the physiological and metabolic measurements performed in 15- to 17-week-old male and female SKO mice. Relative gene expression levels of hepatic metabolic markers in the liver of AAV-aP2 (**B**) and AAV-TBG (**C**) injected mice eight weeks after I.P. injection of $1\times10^{12}$ genome copies. (**D**) Tissue weights of retroperitoneal white adipose tissue (rWAT) from WT, AAV-CMV, AAV-aP2 and AAV-TBG injected mice. Relative gene expression levels of adipocyte markers in gWAT of AAV-aP2 (**E**) and AAV-TBG (**F**) injected mice eight weeks after I.P. injection of $1\times10^{12}$ genome copies. (**G**) Western blot tissue panel analysis (excluding the liver) of human BSCL2 protein levels in AAV-TBG injected mice. All data are biological replicates presented as the mean ± SEM, n = 16 (WT), 5-6 (AAV-GFP), 5 (AAV-aP2) and 5 (AAV-TBG) mice per group for B, C, E and F, n = 44 (WT), 5 (AAV-CMV), 5 (AAV-aP2) and 5 (AAV-TBG) mice per group for D, * $p<0.05$, ** $p<0.01$, *** $p<0.001$ and **** $p<0.0001$ vs WT, ## $p<0.01$, ### $p<0.001$ and #### $p<0.0001$ vs AAV-GFP, ntd = no tissue dissected, $ = not detected, L indicates the presence of a molecular ladder.

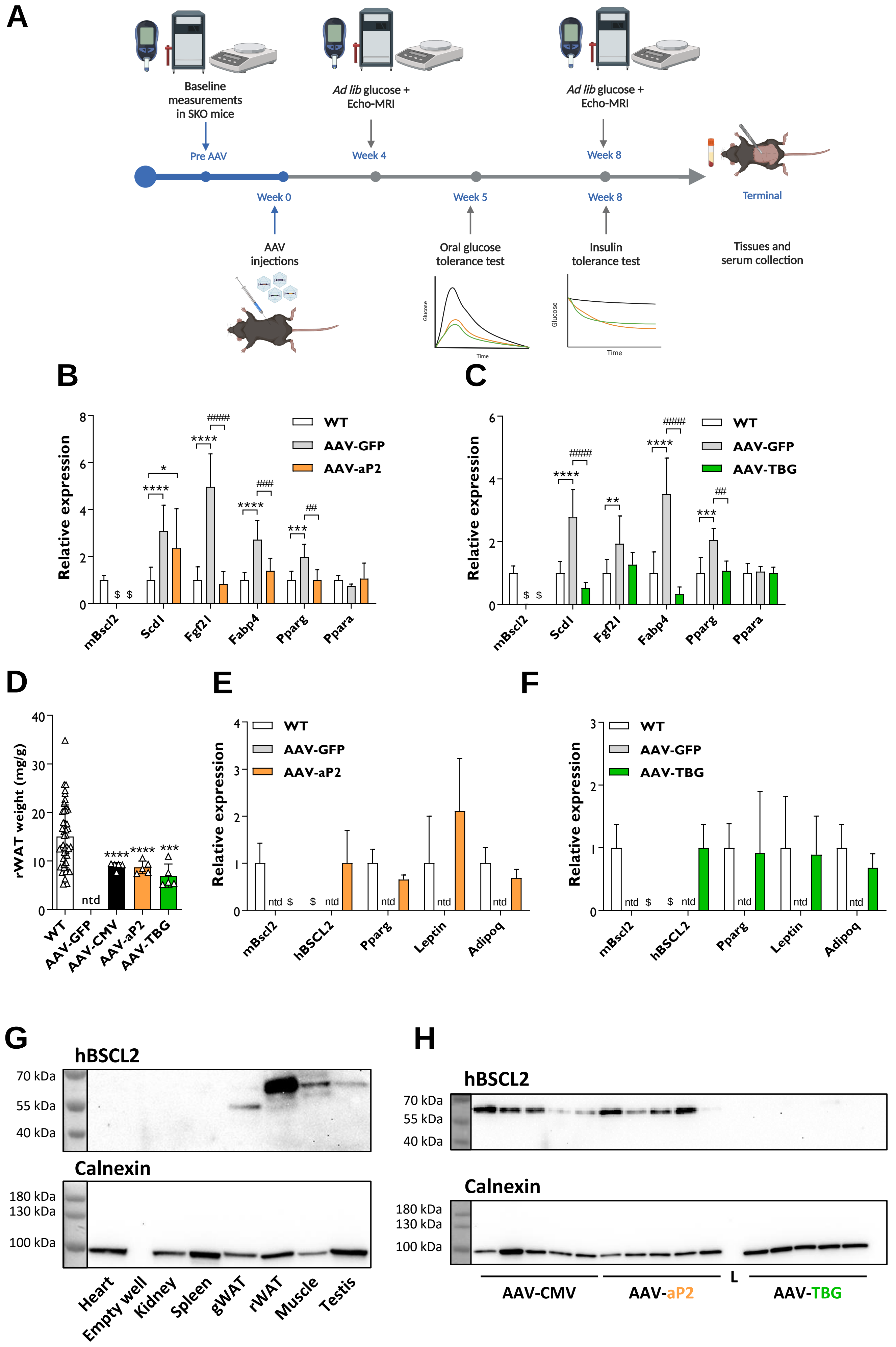
A
Baseline measurements in SKO mice
Pre AAV
Ad lib glucose + Echo-MRI
Week 4
Ad lib glucose + Echo-MRI
Week 8
Week 0
Week 5
Week 8
Terminal
AAV injections
Oral glucose tolerance test
Insulin tolerance test
Tissues and serum collection
Glucose
Time
B
Relative expression
WT
AAV-GFP
AAV-aP2
mBscl2
Scd1
Fgf21
Fabp4
Pparg
Ppara
C
Relative expression
WT
AAV-GFP
AAV-TBG
mBscl2
Scd1
Fgf21
Fabp4
Pparg
Ppara
D
rWAT weight (mg/g)
ntd
WT
AAV-GFP
AAV-CMV
AAV-aP2
AAV-TBG
E
Relative expression
WT
AAV-GFP
AAV-aP2
mBscl2
hBSCL2
Pparg
Leptin
Adipoq
F
Relative expression
WT
AAV-GFP
AAV-TBG
mBscl2
hBSCL2
Pparg
Leptin
Adipoq
G
hBSCL2
70 kDa
55 kDa
40 kDa
Calnexin
180 kDa
130 kDa
100 kDa
Heart
Empty well
Kidney
Spleen
gWAT
rWAT
Muscle
Testis
H
hBSCL2
70 kDa
55 kDa
40 kDa
Calnexin
180 kDa
130 kDa
100 kDa
AAV-CMV
AAV-aP2
L
AAV-TBG
